# Sex, Stress and the Heart: Long-term Cardiovascular Effects of Embryonic Metabolic Disruption

**DOI:** 10.1101/2025.03.21.644511

**Authors:** J.J. Lees, B. Bicici, S. Berglund, K. Smith, G. Galli, J. Altimiras, C. Guerrero-Bosagna

## Abstract

Adverse conditions within the embryonic environment can alter embryogenesis, programming systemic physiological changes that may manifest as disease states in adult life. The process of developmental programming represents an important factor underlying cardiometabolic diseases, many of which are leading causes of death globally. Importantly, there is evidence that males are less tolerant to certain environmental perturbations during embryogenesis, mirrored by sex differences in the incidence of certain cardiometabolic diseases. Understanding sex differences in programmed responses in mammalian models is complicated by maternal compensation and placental factors. Avian models offer a valuable comparable system in which such effects are not present. Here, we investigate the influence of developmental hypoxia and hypothermia in programming cardiovascular structure and function in the domestic chicken (*Gallus gallus domesticus*). In agreement to mammalian studies, adult males but not females show pathological mitochondrial morphology and respiratory capacity, ventricular hypertrophy and reduced body weight programmed by embryonic hypothermia and hypoxia. These data not only represent novel findings in birds but demonstrate the utility of the avian model for understanding sex differences in prenatal stress responses, revealing common responses among endothermic amniotes.

## Introduction

Embryonic development is a period during which organisms are highly susceptible to the deleterious impacts of environmental stressors. Exposure to both acute and chronic environmental or chemical exposures during this period drive physiological responses that impact embryonic development ^1^. Subsequently, through ‘developmental programming’, embryonic gene-environment interactions in response to developmental stress may manifest as persistent pathophysiological phenotypes or increased susceptibility to environmental perturbations in adults long after the stressor is removed ^2^. Importantly, sex differences are often observed in the physiological responses to developmental stress ^3^. Of these responses, cardiovascular function is strongly influenced by the neonatal environment, given the central role of the circulatory system in supplying the metabolic processes required for development^4^. The concept that prenatal environmental conditions shape long-term cardiovascular risk is well established in human epidemiology, where low birth weight—a surrogate marker for intrauterine stress—is associated with an elevated incidence of cardiovascular disease in adulthood ^5^. Additionally, sex differences in the developmental programming of cardiovascular disease are well-documented, with men generally exhibiting higher rates of cardiovascular morbidity and mortality compared to women, a disparity attributed to both genetic and developmental factors ^6^. Given that cardiovascular diseases represent the primary causes of death worldwide, with key differences in susceptibility between the sexes ^7^, investigating the interaction between sex and developmental stress is central to our understanding of cardiometabolic disease ^8^. However, despite the predominance of cardiovascular disease later in life, developmental programming studies often focus upon the embryonic and early postnatal development effects. More data are therefore required integrating multiple biological and temporal scales, whilst considering the influence of sex.

Oxygen availability is an important source of environmental stress in avian embryos and has overarching impacts upon development. Depending on the species, timing and length of exposure, hypoxia and hypothermia drive cardiovascular reprogramming at multiple hierarchical levels in a sex-specific manner. In mammalian models, adult male rats exposed to developmental maternal hypoxia display growth restriction, cardiomyopathy ^9^, diastolic dysfunction ^10^ as well as increased susceptibility to ischaemia and reperfusion injury ^11^, increased heart/body weight ratio ^12^, increased myogenic tone ^13^ and fibrosis ^14,15^. Concurrent with these morphological and functional differences, cardiac mitochondria in mammals also respond to developmental hypoxia in a sex-specific manner. For example, adult male mice and fetal male rats from hypoxic pregnancies show increased H_2_O_2_ production and a lower mitochondrial respiratory capacity than equivalent females ^16,17^ similar to findings in Guinea pigs ^18,19^.

Investigating the developmental effects of environmental stressors in isolation is challenging in mammalian models due to confounding maternal and placental effects. Such effects are not present in birds, making them an excellent model for embryonic stress in which the exposure of the embryo can be more precisely controlled. As a result, much has been done to investigate the embryonic effects of developmental hypoxia and hypothermia in birds. Depending upon the developmental window of exposure, prenatal hypoxia has been shown to affect embryonic development by blunting the development of thermogenesis ^20^ and produces structural changes in the eye and beak ^21^ as well as the liver ^22^. Cardiovascular structure and function are also altered in embryos incubated under hypoxia, which results in reduced ventricular pressure ^23^, elevated heart weight ^24^, reduced cardiomyocyte number ^25^ and sensitization of β-adrenoceptors ^22^ associated with postnatal cardiac contractile dysfunction^26^. Prior to the acquisition of endothermy, birds are also susceptible to fluctuations in temperature. Chicken embryos are poikilothermic during the first 18 days of incubation and so temperature influences metabolism, nutrient usage and embryonic development. Hypothermia, in turn, is shown to produce delayed hatching and chicks with low hatching weight ^27^, delayed hatching, growth retardation and embryonic cardiac hyperplasia in chickens ^28,29^.

Despite the extensive demonstration of embryonic and early life effects of incubation stress, it remains important to connect embryonic conditions with the adult cardiac phenotype, given that the severe consequences of cardiovascular dysfunction often manifest later in life. However, data regarding the life-long programmed effects linking cardiometabolic disease to programming during development have received relatively less attention in birds in comparison to mammals, particularly when considering sex differences. Furthermore, disparities between existing studies regarding levels of exposure, timing, age of sampling and the biological response of interest make comparison difficult. Taken together however, data indicate that male chicken hearts may be more susceptible to the long-term effects of hypoxia during development. Experimental hypoxia of 13.5 % to 15% O_2_ results in left ventricular wall thickening in adult hens ^30^ and altered arterial reactivity in adult chickens ^31^. Similarly, high altitude incubation results in elevated heart weight and pulmonary artery diameter in 6 month old birds ^32^. Studies investigating the mitochondrial implications of hypoxic incubations are sparse, with only a single study demonstrating reduced complex IV oxygen flux, in one day old chicks incubated under 15% oxygen ^33^. The longer-lasting implications of developmental hypothermia have received relatively little attention in comparison to hypoxia studies. Broiler chickens show reduced hatch weight which is retained at 6 weeks post hatch resulting from hypothermic (36.6°C incubation) ^34^ and show evidence of higher relative heart weights as well as reduced incidences of ascites during growth ^35^. In terms of mitochondrial metabolism, long-lasting elevated mitochondrial aerobic metabolism was observed in 60 day old Japanese quail as a result of elevated incubation temperature although sex differences in this parameter were not reported ^36^.

Here we determine the extent to which developmental stress programmes long-term cardiovascular changes in male and female chickens. For this, we use a combination of hypoxic and hypothermic incubations to then evaluate the cardiometabolic effects over multiple biological levels in adults reared under normal environmental conditions. For the first time in birds, we demonstrate significant sex differences over multiple hierarchical levels from mitochondrial protein levels and respiration to whole animal metabolism. The results indicate that, independent of maternal effects, alterations to the embryonic environment programme lifelong cardiometabolic alterations which manifest differently dependent upon sex.

## Materials and Methods

### Ethics

All experiments were performed under the ethical permission number 5.2.18-11569/2020, granted by the Uppsala Animal Experimentation Ethics Committee. Birds were housed in industry-standard conditions during the experiments and euthanized following …Swedish legislation.

### Incubation and treatment groups

Eggs of Bovans Robust hybrid egg-laying chickens (*Gallus gallus domesticus)* were obtained from an industrial supplier (SwedFarm AB). During the first 17 days of incubation, eggs were placed under 4 different oxygen and temperature conditions: control (under atmospheric O_2_ ∼ 21% and 37.8°C), hypothermic (under atmospheric O_2_ ∼ 21% and 35.8°C), hypoxic (under 18% O_2_ and 37.8°C) and hypoxic hypothermic (‘double treatment’ under 18% O_2_ and 35.8°C). These conditions were relatively mild compared to previous developmental studies in order to ensure hatchability of eggs and survival of birds to sexual maturity. Hypoxia was maintained by nitrogen flow, regulated using a Roxy-4 gas regulator (Sable systems International, Las Vegas USA). At day 18, eggs from all treatment groups were then transferred to a hatcher at 37.5 °C, 65% relative humidity and normoxic conditions for hatching.

### Animals and husbandry

Upon hatching, birds were wing-tagged, weighed and transferred to indoor cages. Control (N = 39, 19 male and 20 female), Hypoxic (N = 40, 18 male and 22 female), Hypothermic (N = 24, 12 male and 12 female) and Double treatment (N = 15, 7 male and 8 female) individuals were then reared with *ad libitum* access to food and water under the lighting and temperature recommendations for the breed. Birds were provided with perches and were weighed at regular intervals throughout the experiment.

### Whole body measures and sample collection

Birds were killed by stunning followed by decapitation at between days 256-275 during which all sample collection and subsequent cardiac measures took place. Blood samples for telomere analysis were collected via the ulnar vein. Blood was collected in in sodium-heparin capillary tubes, centrifuged at 600g for 5 minutes and the red blood cells retained for DNA extraction. For dissections and heart sampling, animals were immediately dissected and livers and hearts removed for weighing. Whole livers were weighed after blotting to remove superficial water. Blood was removed from the chambers of the heart manually before blotting to remove superficial water before weighing. The atria were then removed before the left and right ventricles were separated and weighed, each retaining approximately half of the septum. Tissue from the apex of the left ventricle was then removed and divided into separate samples for mitochondrial respirometric analysis (performed on fresh tissue), DNA and protein extraction (frozen samples) and fixation for electron microscopy. Finally, the tarsus length was measured with digital callipers as a proxy for size.

### Respirometry – Egg and whole animal

To determine the metabolic effects of our treatments, eggs’ metabolism was measured at day 17, using flow through respirometry in a pull-mode configuration. Eggs were placed into custom sealed metabolic chambers (200 ml) within their respective incubators to ensure maintenance of incubation temperature. Similarly, incurrent air was drawn from the eggs’ incubators (at a flow rate of 200 ml/min) to maintain oxygen and humidity levels consistent with treatment conditions. Gas was first scrubbed of water (using calcium chloride) and carbon dioxide (using sodalime) prior to measurement of oxygen concentration using a FoxBox field gas analysis system (Sable systems International, Las Vegas USA), which also served as the pump pulling air through the chambers. Eggs were not sexed prior to analysis.

As well as measuring the direct effects of our treatments upon metabolism, we quantified the longer-term effects of the incubation treatment upon the metabolism of the birds post hatch. To do this, whole animal metabolism was measured, again using pull, flow through respirometry. Birds (age 64-77 days) were placed in a sealed chamber (15 litre volume) within a temperature-controlled incubator shortly after the start of the dark period of their light:dark cycle. Incurrent oxygen was pushed through each chamber (at a flow rate of 2500 ml/min) and the 4 chambers were multiplexed with an RM-8 respirometry flow multiplexer (Sable systems International, Las Vegas USA). Chamber oxygen was then subsampled at 500 ml/min and scrubbed of water (Using calcium chloride) and carbon dioxide (using sodalime) prior to measurement of oxygen using a FoxBox field gas analysis system (Sable systems International, Las Vegas USA). Oxygen from the incubator was measured between adjacent chamber measurements to create a reliable continuous oxygen baseline. Birds were measured for an average of 3 hours and metabolic measures taken from the last hour of the period to ensure all birds were in a resting state and fasted for at least 2 hours during measurement. Calculation of 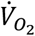 was performed using the equation 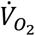, where Fr_i_ is the incurrent flow rate, F_i_O_2_ is the fractional oxygen concentration of the incurrent air and F_e_O_2_ is the fractional oxygen concentration of the excurrent air from the chamber ^37^.

### Mitochondrial respirometry

To isolate the mitochondria, ventricular tissue was minced in ice cold homogenization buffer (MOPS 5mM, NaEDTA 2mM, sucrose 70mM, Mannitol 220 mM, BSA 0.1%). Trypsin (100,000 BAEE units) was added and the tissue was left for 15 minutes before deactivation of trypsin with trypsin inhibitor (6.5 mg, equivalent to inhibition of 100,000 units of trypsin) followed by transfer to a 12 ml glass mortar. Tissue was homogenized on ice using four passes of a loose-fitting pestle at 120 rpm. Mitochondria were separated from cell fractions through centrifugation at 600g for 5 minutes at 4°C. The supernatant was then filtered through gauze and centrifuged at 8500 g for 10 minutes. The mitochondrial pellet was then resuspended in fresh homogenization buffer and centrifuged at 8500 g for 10 minutes. The final pellet was resuspended in 20 μl of homogenization buffer without BSA. The mitochondrial suspension was kept on ice prior to the Oroboros respiration assay and an aliquot was frozen for subsequent Bradford assay of protein content to which all respirometric measures were normalised.

Mitochondrial respiration was measured using an Oroboros Oxygraph 2-k high-resolution respirometry system (Oroboros Instruments, Innsbruck, Austria). Oxygen electrodes were calibrated daily with air-saturated MiRO5 respiration solution (in mM: 0.5 EGTA, 1.4 MgCl2, 20 taurine, 10 KH2PO4, 20 HEPES, 1%BSA, 60 K-MES, 110 sucrose, pH 7.1, adjusted with 5 N KOH). Zero calibrations and O_2_ response calibrations were achieved by injecting titrating dithionite into the experimental chambers. During experiments, two identical respiration chambers (chamber A and chamber B) held at the same temperature were run in parallel. Isolated mitochondria (2-10 microliters of 1.49-10.75 mg protein/ml) were added to each chamber containing 2 ml of respiration medium. All measurements of respiration rates were carried out at adult chicken body temperature (41°C) and with randomisation of treatments and chamber ensuring no chamber bias in the data.

In order to quantify mitochondrial function, we applied a SUIT protocol ^38^. First, complex one substrates pyruvate (5 mM), malate (2 mM) and glutamate (10 mM) were added to achieve Leak respiratory state with complex-I (CI) substrates in the absence of adenylates (Leak_N,CI_). Once a stable oxygen trace was attained, saturating ADP (0.5 mM) was added to generate oxidative phosphorylation with CI substrates (OXPHOS_CI_). Once ADP had been completely phosphorylated to ATP, isolated mitochondria entered state-IV respiration, the Leak respiratory state in the presence of adenylates with CI substrates (Leak_T,CI_). After establishing stable Leak_T,CI_, succinate (10 mM) was then added to assess the contribution of complex-II (CII) substrates to Leak respiration (Leak_T,CI+CII_), and ADP was added again to assess OXPHOS with CI and CII substrates (OXPHOS_CI+CII_). To assess maximal electron transport capacity of the electron transport chain (ET) with CI and CII substrates (ET_CI+CII_), carbonyl cyanide-*4*-(trifluoromethoxy)phenylhydrazone (FCCP) was titrated to a final concentration of 0.1–0.3 μM in order to uncouple mitochondria. Next, the CI inhibitor rotenone (0.5 μM) was added to assess ET_CII_, with CII substrates only. To block the electron transport chain and assess residual non-mitochondrial O_2_ consumption (ROX), the complex-III (CIII) inhibitor antimycin-A (2.5 μM) was added. To assess complex-IV (CIV) activity in isolation, the electron donor N, N, N’,N’-tetramethyl-p-phenylenediamine (TMPD; 0.5 mM) was added in combination with ascorbate (2 mM) to avoid autooxidation of TMPD. Lastly, the CIV inhibitor sodium azide (50 mM) was added to assess background non-mitochondrial O_2_ consumption from the addition of TMPD. To estimate mitochondrial efficiency, the respiratory-control ratio (RCR) was calculated as OXPHOS_CI_/Leak_T,CI_.

### TEM Mitochondrial imaging and analysis

The samples were fixed in 2.5% Glutaraldehyde (Ted Pella) + 1% Paraformaldehyde (Merck) in 0.1 M Phosphate buffer (PB) pH 7.4 and stored at 4°C until further processed. Samples were rinsed with 0.1 M PB for 10 min, after which they were incubated for one hour in 1% osmium tetroxide (TAAB) in 0.1 M PB. After rinsing in 0.1 M PB, samples were dehydrated using increasing concentrations of ethanol (50%, 70%, 95% and 99.9%) for 10 minutes each step, followed by 5 min incubation in propylene oxide (TAAB). The tissue samples were then placed in a mixture of Epon Resin (Ted Pella) and propylene oxide (1:1) for one hour, followed by 100% resin and left overnight. Subsequently, the samples were embedded in capsules in newly prepared Epon resin, left for 1-2 h and then polymerized at 60°C for 48 h.

The specimens were cut into semi thin sections (1-2 μm), stained in Toluidine Blue and examined in a light microscope. The block was trimmed, ultrathin sections (60-70 nm) were cut in an EM UC7 Ultramicrotome (Leica) and placed on a grid. The sections were subsequently contrasted with 5 % uranyl acetate and Reynold’s lead citrate and visualized with Tecnai™ G2 Spirit BioTwin transmission electron microscope (Thermo Fisher/FEI) at 80 kV with an ORIUS SC200 CCD camera and Gatan Digital Micrograph software at 11500x magnification (both from Gatan Inc.). Sample ID was not known at the time of imaging in order to keep the image analysis blind and images were taken from different parts of the sample, chosen at random from the imaging grid.

Morphological analysis of TEM images (1 image per sample) was performed using ImageJ (NIH) software. For each image, the perimeters of all individual mitochondria (average ± stdev = 10 ± 3) were then manually traced using the freehand tracing tool. Average mitochondrial area and aspect ratio (ratio of the major axis to the minor axis) was then calculated for each individual and compared between treatment groups using the default imageJ measurement tool.

### Protein expression levels

To investigate whether functional differences in the mitochondrial electron transport chain were the result of altered protein expression, western blot was used to characterize the levels of complexes I, II, IV and V. Complex III was not measured due to lack of antibody binding to avian samples. Whole homogenates (∼50 mg) were prepared in RIPA buffer containing protease and phosphatase inhibitor cocktail (Sigma, PPC1010) and protein concentration was measured (DC Protein Assay, BioRad, UK). Proteins (10 μg per sample) were separated by PAGE, transferred to nitrocellulose membranes, blocked with 5% milk in TBS-T, and incubated with a primary antibody cocktail (Abcam-ab110413, 1:1000) and an IRDye® 800CW IgG-specific secondary antibody (Licor, 1:20,000). Detection was performed via infrared fluorescence (Licor, UK).

Due to potential variability in housekeeping proteins, total protein staining (REVERT, Licor, UK) alongside an internal control were used for normalization, following Li et al. ^39^. Blots were performed in triplicate on separate occasions and averaged.

The antibody detected four bands, corresponding to the electron transport chain complex subunits: 8 of complex I (NDUFB8), B of complex II (SDHB), 1 of complex IV (MTCO1) and alpha of complex V (ATP5A). Band intensities were normalized to total protein and the internal control. Analysis was performed using Image Studio v5.2.

### Telomere length determination

In order to estimate the rate of aging in the birds the rate of telomere loss was estimated in DNA from red blood cells. DNA was extracted from red blood cells using a standard protocol. Briefly, red blood cells were incubated with DNA lysis buffer (1M trisHCL (pH8), 0.5M EDTA, 10%SDS) at 65ºC for 15 minutes before adding proteinase K and further incubating for a further hour at 55 ºC. After protein precipitation and centrifugation at 12,000 x g for 30 minutes, the DNA was precipitated from the supernatant using isopropanol and washed with 70% ethanol. Telomere abundance was then quantified in the DNA samples using the qPCR method of Criscuolo, et al. ^40^ Primers consisted of: Tel1b (5’CGGTTTGTTTGGGTTTGGGTTTGGGTTTGGGTTTGGGTT-3’) and Tel2b (5’-GGCTTGCCTTACCCTTACCCTTACCCTTACCCTTACCCT-3) and GAPDH (FW-GTCAAGGCTGAGAACGGGAA, RV-GCCCATTTGATGTTGCTGGG). PCR conditions were 10 min at 95°C followed by 40 cycles of 1 min at 56°C and 1 min at 95°C. 2-ΔΔCT values for day 246 were subtracted from those at day 49 to establish a relative decrease in telomere abundance.

### Statistics

Prior to statistical analysis, an outlier test (median absolute deviation test) was run to identify within treatment group outliers and significant outliers were removed. Treatment effects were then determined using a Kruskall-Wallis test and where significant treatment effects were observed a Dunn’s post-hoc test with Benjamini-Hochberg correction was performed to determine differences between treatments. Given the established sexual dimorphism in this species, statistical analyses were conducted separately for males and females. This approach allowed for the identification of sex-specific treatment effects without conflating potential differences in baseline values or variance structures between sexes. All statistics and plotting were performed in R studio (Version 2022.12.1+366, R version 4.4.0 (2024-04-24)). A significance threshold of α = 0.05 was used to determine statistically significant differences between groups for subsequent analyses.

## Results

In this study we investigated the long-term programmed effects of embryonic stress in the form of incubation under hypothermic, hypoxic and double treatment. In comparison to controls, these developmental metabolic stressors induced distinct sex-specific effects on cardiovascular structure and function over multiple levels of organisation.

### Respirometry – Egg and whole animal

To assess the impact of our treatments, it was important to quantify the metabolism of day 18 embryos under their treatment conditions. The whole egg oxygen consumption was affected by the incubation treatment (Figure 1, table 1, supplementary table 1). A *post hoc* test revealed that although all three treatments had 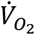 below that of controls, this reduction was only significant in hypothermic and double treatment embryos.

**Table 1:** Summary statistics for whole animal responses of adult birds to incubation under hypoxia, hypothermia and double treatment. X^2^ values, degrees of freedom (df) and p values represent those from Kruskall-Wallis tests for treatment effects. Significant values (bold) represent significant treatment effects within the sex group, with level of significance indicated by stars.

| Measure | Sex | $\chi^2$ | df | p | |
| --- | --- | --- | --- | --- | --- |
| Egg VO2 | NA | **16.7** | **3** | **<0.001** | *** |
| VO2 day 64-77 | Male | 2.4 | 3 | 0.5 |  |
|  | Female | 1.2 | 3 | 0.8 |  |
| Hatch weight (g) | Male | 3.7 | 3 | 0.3 |  |
|  | **Female** | **14.8** | **3** | **<0.01** | ** |
| Weight day 21 (g) | **Male** | **29.5** | **3** | **<0.001** | *** |
|  | **Female** | **31.7** | **3** | **<0.001** | *** |
| Weight day 49 (g) | **Male** | **24.7** | **3** | **<0.001** | *** |
|  | **Female** | **23.1** | **3** | **<0.001** | *** |
| Weight day 120 (g) | Male | 0.26 | 3 | 1 |  |
|  | Female | 4.6 | 3 | 0.2 |  |
| Weight day 256 (g) | **Male** | **10.5** | **3** | **<0.05** | * |
|  | Female | 0.34 | 3 | 0.95 |  |
| Tarsus length day 256 (cm) | **Male** | **17.2** | **3** | **<0.001** | *** |
|  | **Female** | **17.5** | **3** | **<0.001** | *** |
| Egg number per female | NA | 29.3 | 3 | **<0.001** | *** |
| Egg weight per female (g) | NA | 13 | 3 | **<0.01** | ** |
| Heart weight (% body weight) day 256 | **Male** | **8.4** | **3** | **<0.05** | * |
|  | **Female** | **11.6** | **3** | **<0.01** | ** |
| Liver weight (% body weight) day 256 | Male | 2.4 | 3 | 0.5 |  |
|  | Female | 2.1 | 3 | 0.6 |  |
| Left ventricle weight (% heart weight) day 256 | **Male** | **11.4** | **3** | **<0.01** | ** |
|  | Female | 3.3 | 3 | 0.3 |  |
| Right ventricle weight (% heart weight) day 256 | **Male** | **10.3** | **3** | **<0.05** | * |
|  | Female | 6.7 | 3 | 0.08 |  |
| Telomere loss day 49-246 | Male | 9 | 3 | **<0.05** | * |
|  | Female | 2.6 | 3 | 0.5 |  |

**Figure 1:**
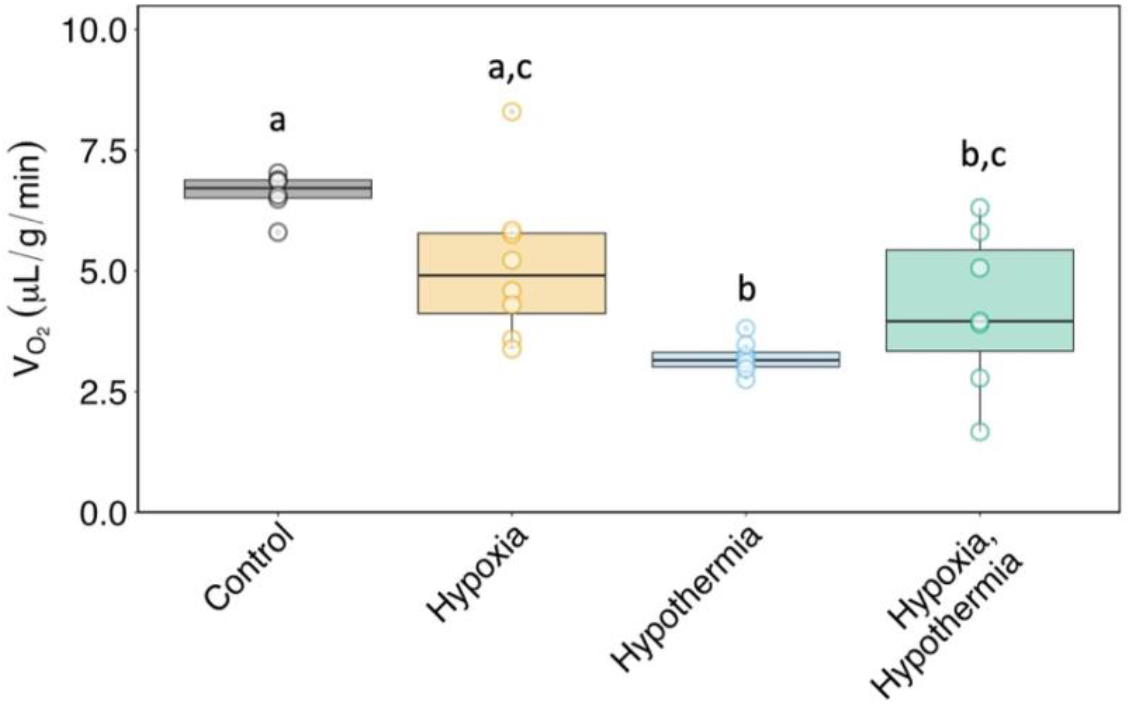
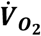 (μl/g/min) of day 18 embryos within their eggs whilst exposed to treatment incubation conditions: control (grey), hypoxic (yellow), hypothermic (blue) and double treatment (green) incubation. Different lower-case letters indicate significant differences between treatment groups /Kruskall-Wallis test, posthoc Dunn test).

To determine if treatments had a long-lasting impact on metabolic rate, resting 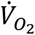 was also measured in adult birds. The oxygen consumption at day 64-77 was not significantly influenced by treatment in either males or females (Figure 2, table 1, supplementary table 1).

**Figure 2:**
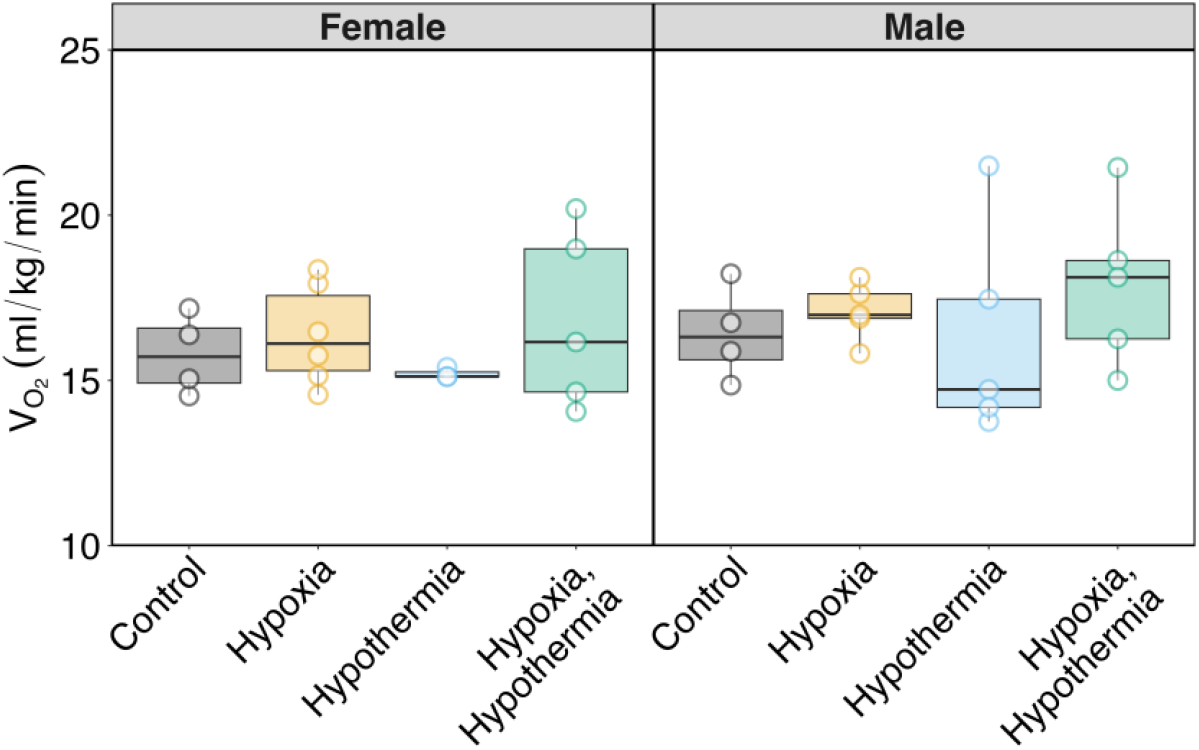
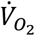 (ml/g/min) of day 64-77 chickens in response to incubation under different temperature and oxygen conditions: control (grey), hypoxic (yellow), hypothermic (blue) and double treatment (green) incubation.

### Growth and adult morphology

To assess if incubation treatments influenced growth, body weight was compared at days 21, 49, 120 and 256. Growth curves differed between males and females, with treatment influencing weight at hatch in females but not males (figure 3, table 1, supplementary table 1). By day 21 and 49, both males and females showed differences in body weight dependent on treatment, however these differences were not apparent by day 256 in females. Male body weight at day 256 was significantly lower in hypothermic birds in comparison to controls. Despite appearing reduced in hypoxic and double treatment males in comparison to controls, this reduction was not significant.

**Figure 3:**
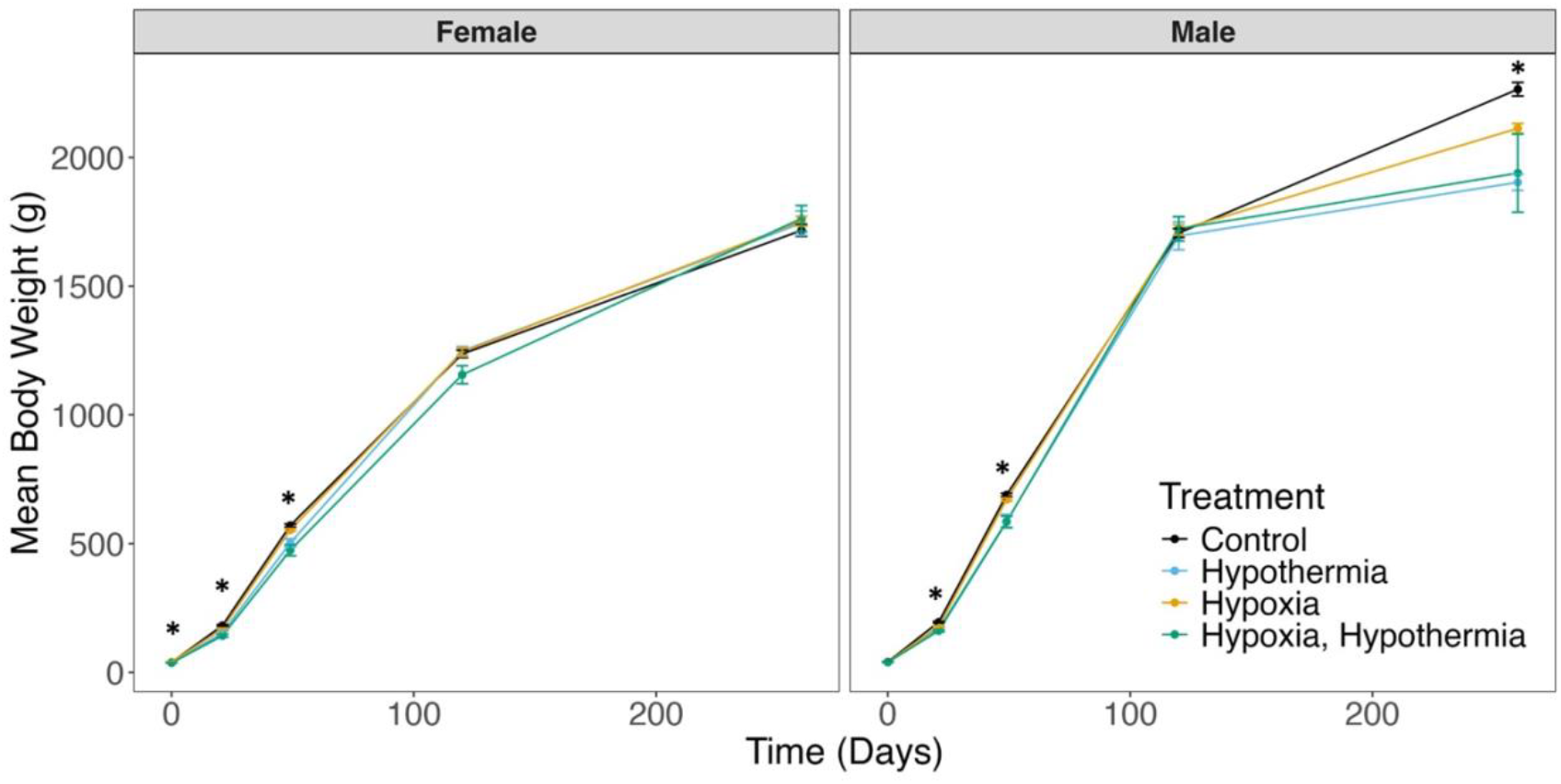
Growth curves of male and female birds in response to incubation under different temperature and oxygen conditions: control (grey), hypoxic (yellow), hypothermic (blue) and double treatment (green) incubation. Stars indicate significant differences between treatment groups within each timepoint (Kruskall-Wallis test, posthoc Dunn test). Error bars represent standard errors of the mean.

As growth can be represented by alterations in both length and volume (weight) parameters, we also compared the tarsus length of individuals as an alternative proxy for size (Figure 4, table 1, supplementary table 1). Tarsus length was decreased by treatment in both males and females. A *post hoc* test showed that both hypothermia and double treatment but not hypoxia reduced tarsus length significantly in males and females. Together, these data show that males experiencing hypothermia stress were both smaller and lighter than control individuals, whereas females retained body weight but were smaller as a result of hypothermia and double treatments.

**Figure 4:**
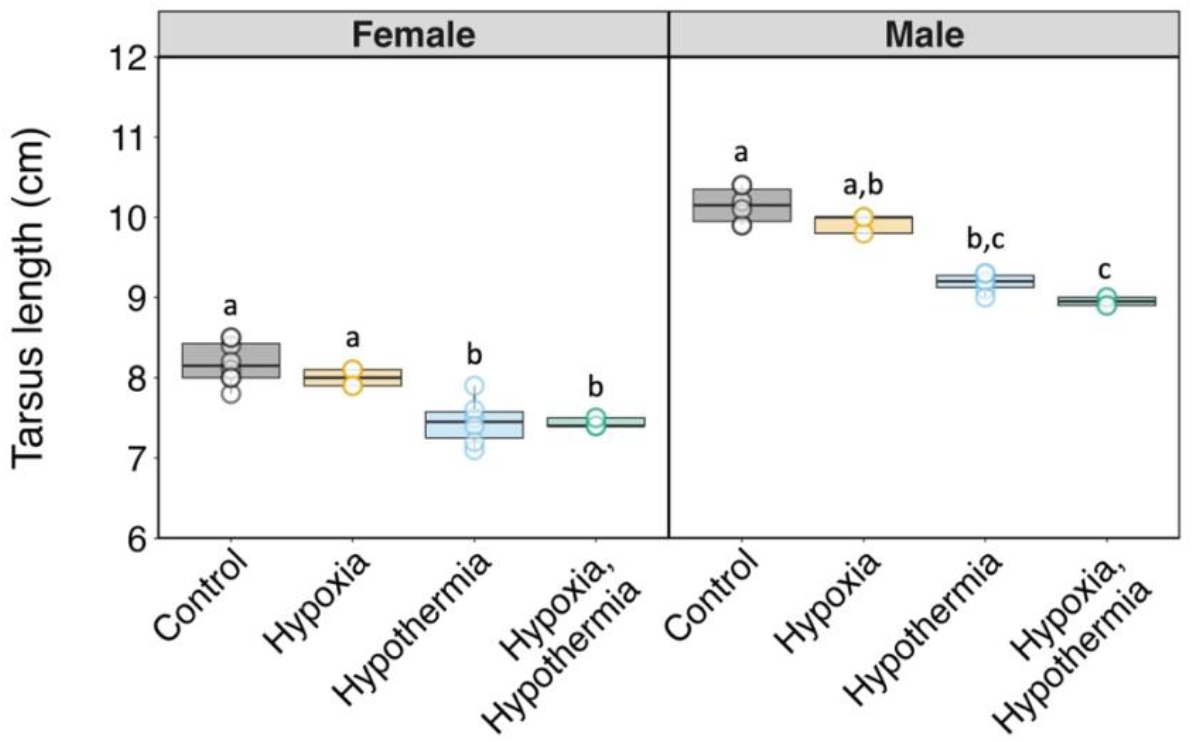
Tarsus lengths (cm) of male and female birds at day 256 in response to incubation under different temperature and oxygen conditions: control (grey), hypoxic (yellow), hypothermic (blue) and double treatment (green) incubation. Different lower-case letters indicate significant differences between treatment groups (Kruskall-Wallis test, posthoc Dunn test).

### Egg laying

Alterations to reproductive output are easily measured in hens as differences in egg numbers and egg weight, which may be representative of physiological stress. Hypothermic and double treatment incubation impacted reproductive output, with females from these groups laying fewer eggs compared to controls (Figure 5, table 1, supplementary table 1). Egg weight was elevated in hypothermia incubated hens, which therefore laid fewer but heavier eggs.

**Figure 5:**
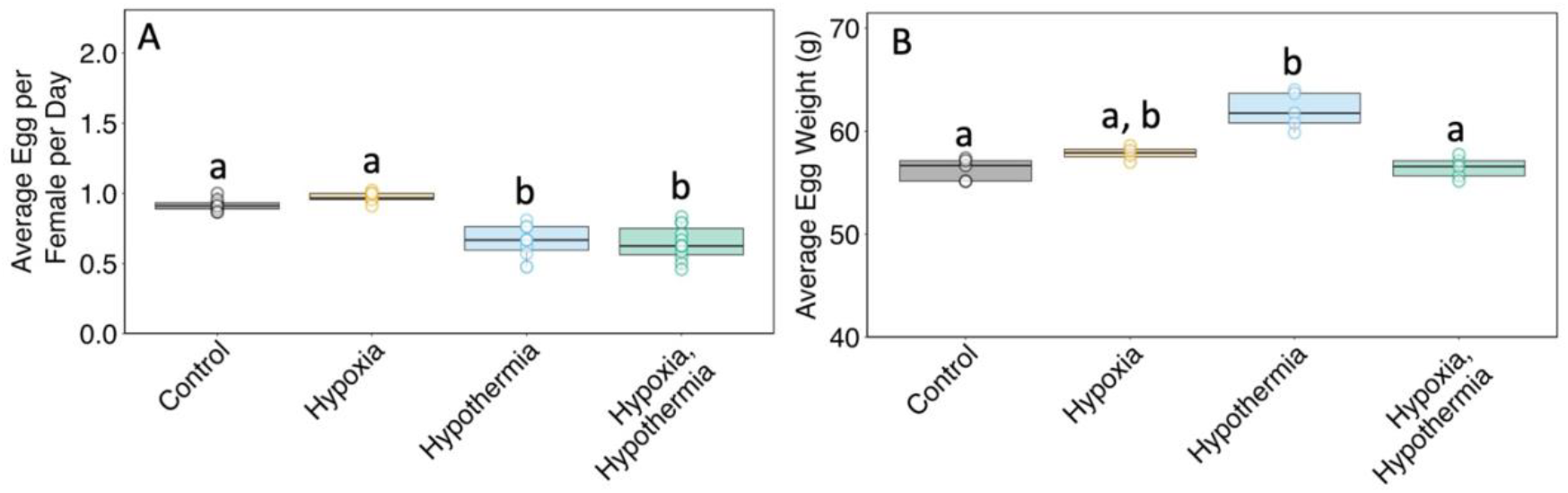
Average number of eggs per day (A) and the average weight per egg (B) laid by females incubated under different temperature and oxygen conditions: control (grey), hypoxic (yellow), hypothermic (blue) and double treatment (green) incubation. Values represent the average of measures taken from all birds between 211 and 243 days old. Different lower-case letters indicate significant differences between treatment groups (Kruskall-Wallis test, posthoc Dunn test).

### Organ size

Because metabolic disruption through incubation has been shown to influence cardiovascular development, organ masses and heart weight parameters were compared in adult birds (Figure 6, table 1, supplementary table 1). Relative liver weight (% of body weight) was unaffected by treatment in males or females, however relative heart weight was influenced by treatment in both males and females. Although relative heart weight (% of body weight) was markedly elevated in hypothermic and double treatment males, this increase was not significant as a result of large variability in the responses. However, relative heart weight in females was significantly higher in the hypothermic group in comparison to controls. Also, only males exhibited significant reductions and increases in left and right ventricular weight (as % heart weight), respectively, as a result of hypothermic and double treatment. Together, these data illustrate that incubation stress not only increases heart weight in both males and females but alters the structure of the ventricles in males.

**Figure 6:**
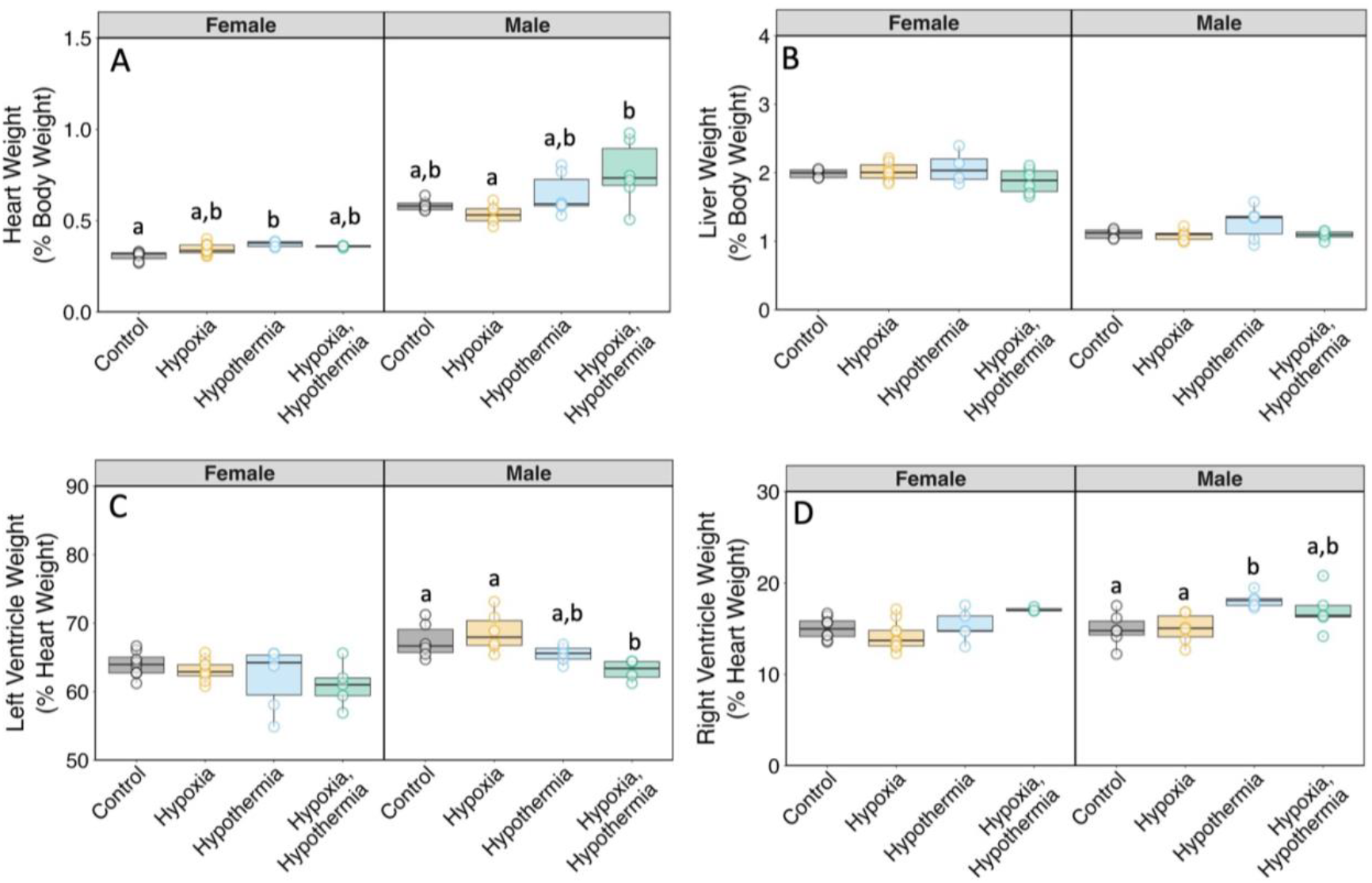
Organ weights of male and female birds at day 256 in response to incubation under different temperature and oxygen conditions: control (grey), hypoxic (yellow), hypothermic (blue) and double treatment (green) incubation. Different lower-case letters indicate significant differences between treatment groups (Kruskall-Wallis test, posthoc Dunn test).

### Telomere length

To investigate the potential for incubation stress to alter the rate of aging of individuals, telomere length loss was compared across individuals (Figure 7, table 1, supplementary table 1). From age 49 to 246 days, telomere length remained unchanged in females, however males from hypothermic incubations showed a reduction in telomere length compared to controls, suggestive of a higher rate of aging in males.

**Figure 7:**
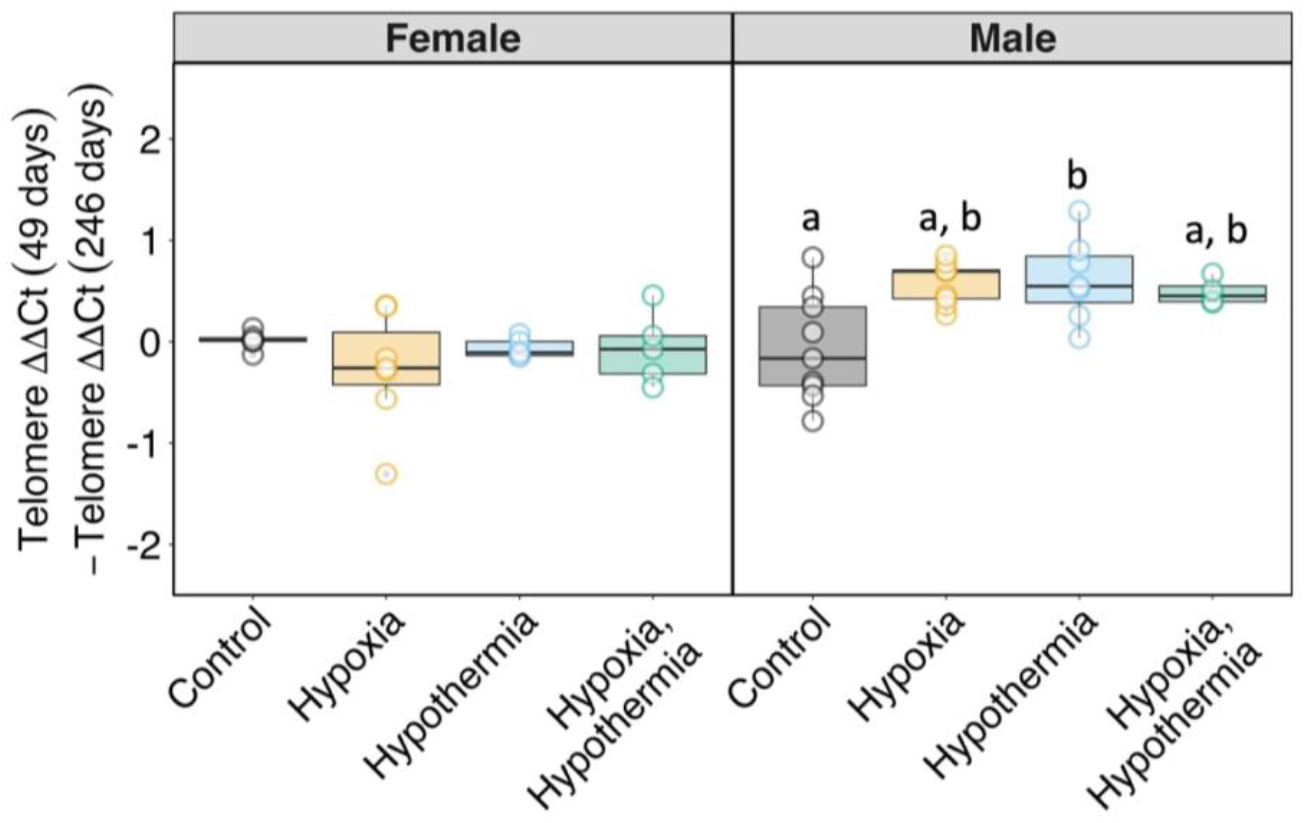
Effects of developmental stress upon telomere loss with age in individuals incubated under different temperature and oxygen conditions: control (grey), hypoxic (yellow), hypothermic (blue) and double treatment (green) incubation. Different lower-case letters indicate significant differences between treatment groups (Kruskall-Wallis test, posthoc Dunn test).

### Mitochondrial morphology

We next investigated whether incubation stress induced long-term alterations to mitochondrial morphology in the cardiomyocytes (Figure 8, table 2, supplementary table 1). Morphological variables such as mitochondrial area and shape represent indicators of both mitochondrial function and oxidative stress in cells. Individual mitochondrial area and aspect ratio was not different in the cardiomyocytes of females from any of the incubation treatments. However, the aspect ratio of male mitochondria was reduced in all treatments, with hypothermic and double treatment incubations being significantly more spherical (aspect ratio closer to 1) than control birds. In order to link mitochondrial morphology with the structural changes in the heart, individual values of mitochondrial area and aspect ratio were regressed against their respective relative heart weight (%). Here, aspect ratio was correlated with relative heart weight in males (ANOVA: F_1_,_15_ = 5.2, r^2^ = 0.21, p < 0.05) but not females, indicating that the mitochondria are more circular in hypertrophic male hearts.

**Table 2:** Summary statistics for cardiomyocyte mitochondrial responses of adult birds to incubation under hypoxia, hypothermia and double treatment. X^2^ values, degrees of freedom (df) and p values represent those from Kruskall-Wallis tests for treatment effects. Significant values (bold) represent significant treatment effects within the sex group, with level of significance indicated by stars.

| Measure | Sex | $\chi^2$ | df | p | |
| --- | --- | --- | --- | --- | --- |
| Mitochondrial area ($\mu\text{m}^2$) day 256 | Male | 2.4 | 3 | 0.5 | |
|  | Female | 3.2 | 3 | 0.4 |  |
| Mitochondrial aspect ratio day 256 | **Male** | **10.6** | **3** | **<0.05** | * |
|  | Female | 0.7 | 3 | 0.9 |  |
| Mitochondrial oxphos CI day 256 | Male | 7.4 | 3 | 0.06 |  |
|  | Female | 5.3 | 3 | 0.1 |  |
| Mitochondrial oxphos CI + CII day 256 | Male | 5.5 | 3 | 0.1 |  |
|  | Female | 4.1 | 3 | 0.3 |  |
| Mitochondrial leak CI day 256 | Male | 1.3 | 3 | 0.7 |  |
|  | Female | 5 | 3 | 0.2 |  |
| Mitochondrial leak CI + CII day 256 | Male | 6.6 | 3 | 0.09 |  |
|  | Female | 1.6 | 3 | 0.7 |  |
| Mitochondrial ET CI + CII day 256 | Male | 4.8 | 3 | 0.2 |  |
|  | Female | 5.4 | 3 | 0.1 |  |
| Mitochondrial ET CII day 256 | Male | 2.4 | 3 | 0.5 |  |
|  | Female | 2.2 | 3 | 0.5 |  |
| Mitochondrial CIV day 256 | **Male** | **11** | **3** | **<0.05** | * |
|  | Female | 2.1 | 3 | 0.5 |  |
| Cardiomyocyte protein CI day 256 | Male | 3.3 | 3 | 0.4 |  |
|  | Female | 4.6 | 3 | 0.2 |  |
| Cardiomyocyte protein CII day 256 | Male | 1.6 | 3 | 0.7 |  |
|  | Female | 4.3 | 3 | 0.2 |  |
| Cardiomyocyte protein CIV day 256 | Male | 3.3 | 3 | 0.4 |  |
|  | Female | 6.9 | 3 | 0.07 |  |
| Cardiomyocyte protein CV day 256 | Male | 1.9 | 3 | 0.6 |  |
|  | **Female** | **11.3** | **3** | **<0.05** | * |

**Figure 8:**
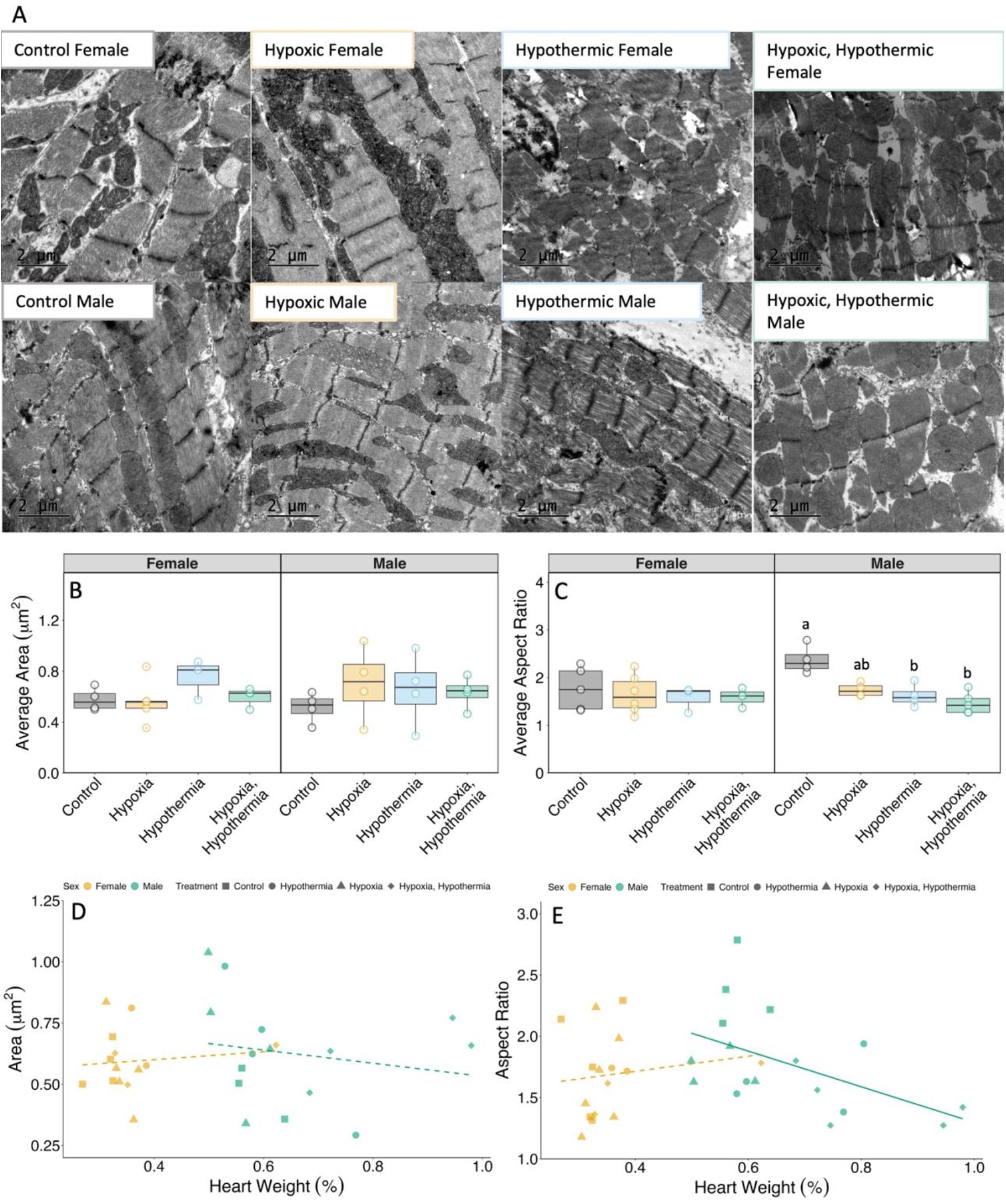
Morphological measures of cardiomyocyte mitochondria taken from transmission electron micrographs. Representative micrographs of ventricular myocardium from male and female birds incubated under different temperature and oxygen conditions (A): control (grey), hypoxic (yellow), hypothermic (blue) and double treatment (green) incubation. Average area (μ^2^) (B) and Aspect ratio (C) of cardiomyocyte mitochondria. Regressions of mitochondrial area (D) and Aspect ratio (E) vs heart weight of the same individuals. Different lower-case letters indicate significant differences between treatment groups (Kruskall-Wallis test, posthoc Dunn test), dashed lines indicate non-significant correlations, bold lines indicate significant correlations.

### Mitochondrial metabolism

To assess whether any cardiovascular effects were associated with alterations to mitochondrial respiration, leak and OXPHOS parameters were compared across individuals (Figure 9, table 2, supplementary table 1). The responses of mitochondrial isolates to all substrates and inhibitors were as expected. None of the mitochondrial respiratory parameters measured was influenced by incubation treatment except for flux through complex IV in males, which was reduced in double treatment males in comparison to hypoxic incubated males. Hypothermic and double treatment males had notably lower (albeit nonsignificant) complex IV flux in comparison to control males. Although not statistically significant, the variability in the oxygen consumption in treated female’s leak_NCI+CII_ state was notably higher than that of males. This apparent difference in variability was not present in the leak_NCI_ state, potentially indicating more variability in the female complex II response to incubation stress.

**Figure 9:**
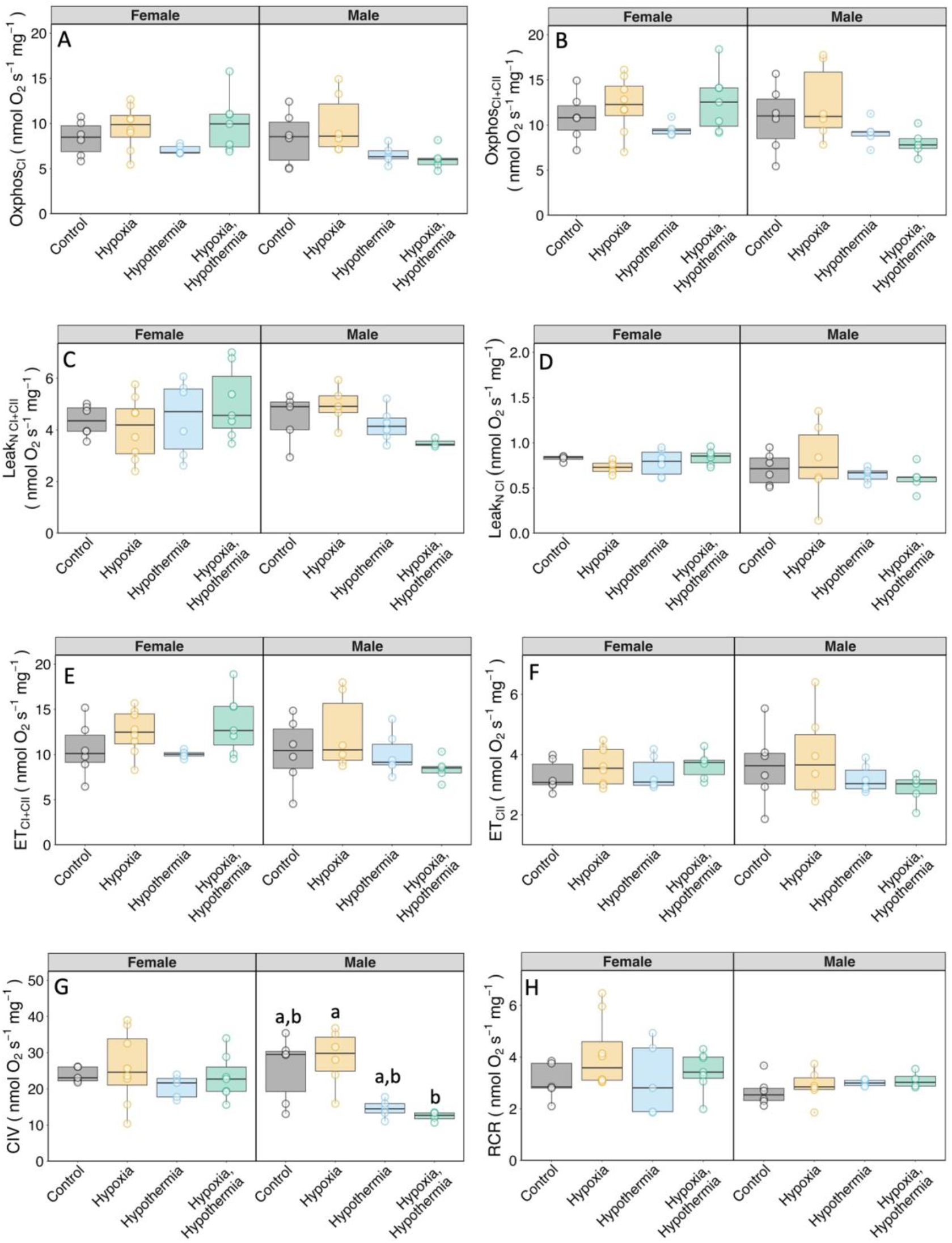
Mitochondrial respiratory parameters of cardiomyocyte mitochondria under different respiratory states. Respiratory parameters of mitochondrial isolates from the left ventricles of males and females incubated under different temperature and oxygen conditions: control (grey), hypoxic (yellow), hypothermic (blue) and double treatment (green) incubation. Parameters include Leak respiratory state with complex-I (Leak_NCI_) and complex-II (Leak_NCI+CII_) substrates in the absence of adenylates, oxidative phosphorylation (OXPHOS) with complex-I (OXPHOS_CI_) and complex-II (OXPHOS_CI+CII_) substrates, maximal electron transport capacity of the electron transport chain (ET) with complex-I (ET_CI_) and complex-II (ET_CII_) substrates, maximal flux through complex IV (CIV) and the respiratory control ratio (RCR). Different lower-case letters indicate significant differences between treatment groups (Kruskall-Wallis test, posthoc Dunn test).

### Mitochondrial electron transport chain complex protein levels

To explore the mechanistic causes of differences in mitochondrial functional, we quantified protein levels of the mitochondrial electron transport chain complexes I, II, IV and V (ATP synthase) (Figure 10, table 2, supplementary table 1). Despite differences in complex IV flux in males, no differences in complex IV protein levels were present. The only significant effects of incubation treatment were upon the levels of ATP synthase (complex V) in females, which were elevated by the hypothermic incubation.

**Figure 10:**
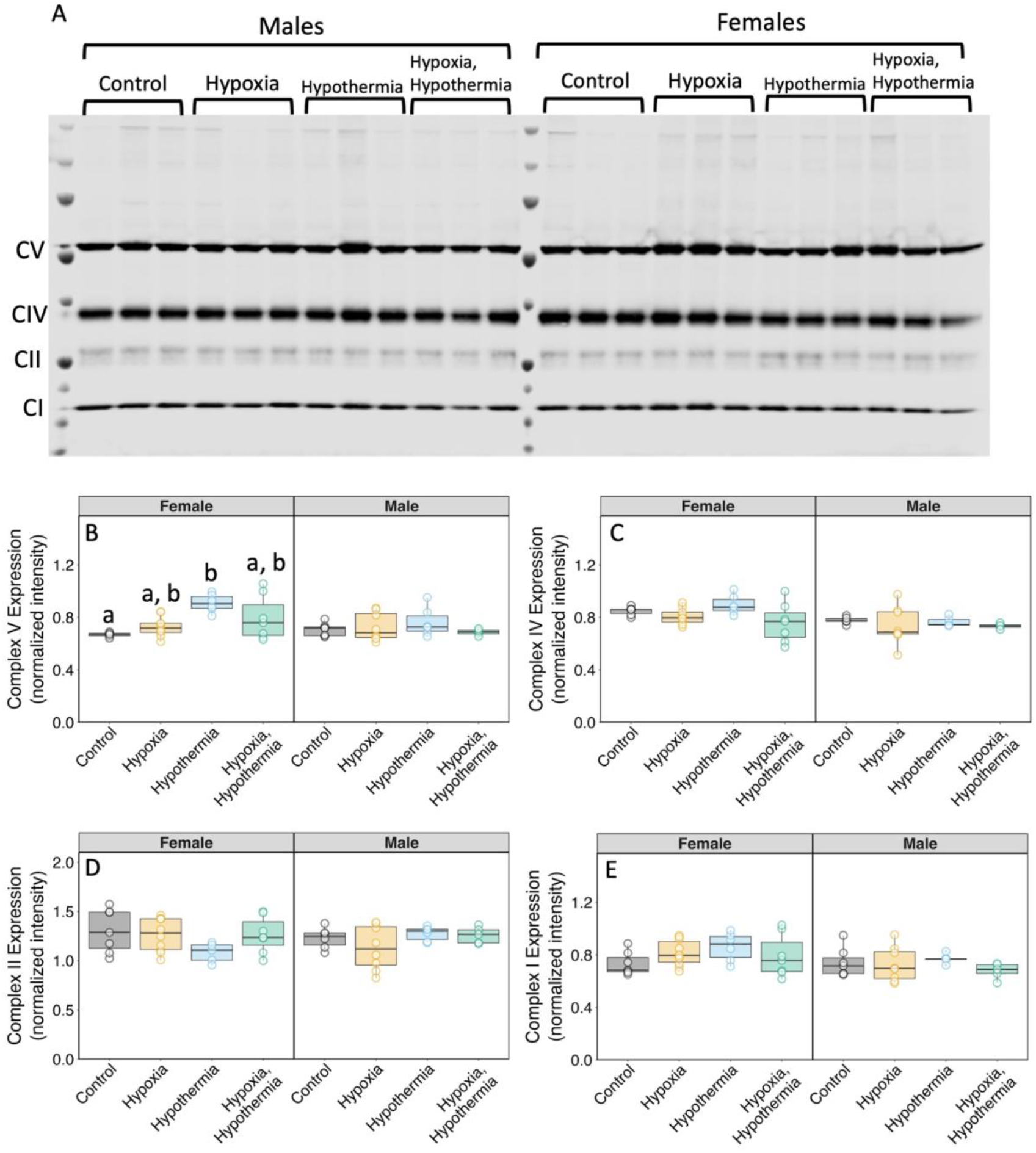
Effects of developmental stress upon the protein expression of respiratory complexes. Relative protein levels were quantified using western blots. The full blot (A) and individual plots of respiratory complex V (CV, B), complex IV (CIV, C), complex II (CII, D) and complex V (CV, E) are shown separately in adult males (M) and females (F) under control (grey), hypoxic (yellow), hypothermic (blue) and double treatment (green) incubation. Different lower-case letters indicate significant differences between treatment groups (Kruskall-Wallis test, posthoc Dunn test)

## Discussion

Here, using an avian model, we quantify long-term developmentally programmed effects over multiple hierarchical levels, spanning protein expression, mitochondrial function, and whole-animal physiology, with a focus on the heart. Across most of the variables measured, hypoxia, hypothermia and double treatment elicited a similar pattern of effects, although significant effects were seen mostly in the adults experiencing hypothermic and double treatment incubations. Overall, these suboptimal developmental environments programmed cardiovascular and metabolic phenotypes in adult birds, underscoring the protracted nature of these developmental effects. Importantly, reductions in temperature and oxygen during development elicited distinct sex-specific effects on cardiovascular structure and cardiomyocyte mitochondrial function. By filling existing gaps in the avian data, these results suggest consistent sex differences in programmed cardiovascular responses across endothermic amniotes. Male, but not female adult birds exhibit reduced body weight, ventricular hypertrophy, reduced mitochondrial bioenergetics and accelerated aging in response to developmental hypoxia and hypothermia, implicating sex-specific pathways in long-term cardiovascular dysfunction. Together, our results reinforce the necessity of integrating sex as a biological variable in developmental programming research as well as the importance of time as a factor in the progression of programmed disease.

Gross morphological variables such as body weight and growth patterns provide insights into the developmental consequences of metabolic stressors. Here, although individuals from both sexes were smaller as adults resulting from hypothermic and double treatment incubations, male and female birds exhibited differences in body weight trajectories. Males retained post-hatch body weight differences from hypothermic incubation whereas females returned to control weight at around 120 days of age. Similar results have been reported in rats in which males and females exposed to fetal hypoxia are similar at four months of age but males have smaller body weight and left ventricular hypertrophy at 12 months ^41^. Such findings underscore the importance of time in developmental programming studies, as early effects may resolve or persist depending on sex-specific compensatory mechanisms, depending on the growth parameter being measured. Interestingly, the catch-up growth in females was not associated with a reduction in telomere aging, as may be expected as a result of the compensation ^42^. However, the observed alterations to egg laying and egg weight after hypothermic incubation suggest that this resilience comes at a cost, influencing reproductive output. Telomere length is a well-established biomarker of biological age and a key factor in determining lifespan. Our findings indicate that males subjected to hypothermic incubation exhibited significantly greater telomere attrition, suggesting an accelerated aging trajectory. This observation agrees with studies in birds showing that early-life stress accelerates telomere shortening, with long-term consequences for survival and lifespan ^43^. More broadly, in humans ^44^, prenatal stress has been linked to increased telomere erosion in males, which has been associated with heightened risks of age-related diseases including metabolic syndrome ^45^ and cardiovascular dysfunction ^46^. Although the mechanisms underlying the sex-specific effects observed were not investigated, more rapid telomere attrition likely reflects the integrated, cumulative effects of altered cardiac and mitochondrial physiology, cellular turnover rates as well as oxidative stress. Future studies should investigate whether the observed telomere attrition translates into reduced longevity and increased disease susceptibility later in life, particularly when subjected to additional environmental stressors. Although not explicitly quantified in this study, it was noted that males from all treatment groups exhibited a higher frequency of malformations in bill and toe morphology, consistent with findings in birds in which beak length is reduced as a result of early hypoxic exposures _21_, which may result from oxidative stress or lack of sufficient perfusion at these sites during development.

Interpreting the cardiovascular effects of developmental stress requires careful differentiation between embryonic, post-hatch, and adult responses. The timing of assessment can significantly impact the observed effects and their interpretation. In embryonic stages, cardiovascular remodelling in response to hypoxia and hypothermia typically manifests as changes in heart mass, ventricular structure, and alterations in the expression of key metabolic regulators ^17,24,25,47^. These early changes set the foundation for long-term cardiovascular function and may serve as predictive markers of later disease risk. Post-hatching, cardiovascular responses can then diverge significantly depending on whether immediate compensatory mechanisms mitigate early structural and functional deficits. For example, embryonic hypoxia has been shown to sensitize β-adrenoceptors, but this effect is reversed postnatally, leading to a desensitization that may compromise long-term cardiac adaptability ^48,49^.

In adults, the long-term effects of developmental stress may therefore become more pronounced as cumulative physiological strain reveals latent vulnerabilities ^5^. Adult birds in our study showed significant gross cardiovascular alterations in response to developmental stress, including increased relative heart weight due to hypothermic treatments in both sexes. However, males exhibited a greater, albeit more variable, increase that was not statistically significant, perhaps due to the observed differences in body weight in males compared to females. In contrast, ventricular weights relative to heart weight were affected only in males, suggesting a sex-specific response in cardiac remodelling. These findings suggest a heightened susceptibility to early-life metabolic insults that may predispose males to long-term cardiovascular dysfunction. The male-biased vulnerability to developmental cardiovascular remodelling aligns with mammalian studies illustrating that male fetuses exhibit heightened susceptibility to hypoxic insults ^50^. Rodent models have demonstrated that prenatal hypoxia predisposes male offspring to diastolic dysfunction, oxidative stress accumulation, and heightened ischemia-reperfusion injury susceptibility relative to females ^10,11^. Similarly, chronic in ovo hypoxia has been shown to induce biventricular cardiac hypertrophy in chicken embryos, alongside reduced pulmonary arterial contractile reactivity ^51^. Interestingly, our data show that hypoxia had no significant effect on heart weight. This is potentially because our hypoxic incubation (18%) was modest compared to that previously shown to affect cardiac development in chickens (13-15% O2) e.g. ^23,24^.

In terms of the mechanisms underlying cardiac hypertrophy, one possibility is the systemic circulation challenges that arise during embryonic development, where stress-induced alterations to vascular resistance, haematocrit and heart rate as a response to systemic hypoxia alter cardiac workload and metabolic demand ^31,52^. The consequences of these developmental alterations may become exaggerated with age, as postnatal cardiovascular remodelling continues in response to increased metabolic demands and changes in hemodynamic load. Additionally, the greater metabolic challenges imposed on male hearts, associated with sexual dimorphism in chickens, may contribute to a heightened susceptibility to long-term cardiac dysfunction compared to females. Conversely, the reduced ventricular effects observed in females align with observations in mammalian studies, where female offspring subjected to prenatal stressors maintain superior cardiovascular adaptability ^6,53^. Interestingly, Lindgren and Altimiras ^22^ show the cardiovascular effects of developmental hypoxia differ dependent on the breed of chicken exposed. Further investigation into the role of different artificially selected production traits upon susceptibility to developmental stress could provide valuable insights into the mechanisms underlying cardiac remodelling and later-life pathology. Potential mechanisms underlying female resilience involve oestrogen-mediated cardioprotection, as oestrogen has been shown to be protective in the programming of hypertension ^8^ in addition to enhancing mitochondrial efficiency, reducing oxidative stress, and promoting vasodilation ^54^. Female mitochondria typically exhibit higher oxidative phosphorylation efficiency and enhanced antioxidant defence mechanisms, which can mitigate the damaging effects of metabolic stress ^54^.

Although gross cardiovascular structure is fundamental to cardiovascular function, the underlying metabolic performance (here reflected in mitochondrial parameters), is equally critical. Hypoxic and double treatment males exhibit a marked reduction in mitochondrial aspect ratio which correlated with relative heart weight increases. More circular mitochondria indicate a shift toward a more fragmented mitochondrial network, a hallmark of mitochondrial dysfunction and compromised oxidative phosphorylation efficiency ^55,56^. Similar reductions in aspect ratio have been demonstrated in Tibetan sheep in response to developmental hypoxia ^57^. Interestingly, such morphological changes have never been investigated in adult birds. In addition to altered morphology, cardiomyocyte mitochondria of hypothermic and double-treatment males show a tendency (non-significant) towards decreased complex IV oxygen flux, a finding congruent with the only comparable study available in one day old chicks incubated under 15% oxygen ^33^. Similarly, in mammalian models, males from hypoxic pregnancies exhibit impaired mitochondrial respiratory capacity in complex II in rats, ^17^), complexes I and IV in mice, ^16^) and complex IV in guinea pigs, ^19^ and rats ^58^. Complex IV, also known as cytochrome c oxidase, plays a critical role as the terminal enzyme of the electron transport chain, where it facilitates the final transfer of electrons to molecular oxygen, allowing for ATP synthesis. A reduction in complex IV activity can therefore lead to diminished oxidative phosphorylation efficiency, electron leakage and increased susceptibility to oxidative stress in male hearts. In mammals and birds, deficiencies in complex IV activity are therefore associated with metabolic disorders and cardiovascular dysfunction ^59-61^.These findings highlight the critical role of mitochondrial respiratory function in developmental and long-term cardiovascular health. One unresolved question, however, is the extent to which the observed alterations to mitochondrial respiration are causative, or the consequence of, the observed ventricular hypertrophy in treated males.Indeed, alterations to myocardial metabolism may not be linked to the development of hypertrophy as seen in female rats ^41^ Rather, these two features may have parallel and synergistic effects on later heart susceptibility to IR injury. Alternatively, males, being more susceptible to the mitochondrial dysfunction caused by hypothermic stress, may have a reduced mitochondrial capacity (specifically at complex IV), which would compromise ATP generation and lead to a compensatory increase in heart mass (ventricular hypertrophy) to maintain cardiac output. In contrast, females may experience less mitochondrial impairment due to greater mitochondrial resilience, possibly driven by oestrogen-mediated protection, and therefore would not develop hypertrophy under the same conditions.

At the level of the mitochondria, females appear to display minimal alterations in response to developmental hypoxia or hypothermia and no alterations in mitochondrial respiration or morphology. However, females demonstrated an upregulation of ATP synthase protein in response to hypothermic incubation, which was not seen in males. This suggests an alternative metabolic adaptation wherein females prioritize ATP production. However, despite alterations in ATP synthase protein levels, no differences were observed in oxidative phosphorylation measurements using Oroboros respirometry. One possible explanation is that ATP synthase activity is regulated post-translationally, meaning increased protein expression does not necessarily equate to increased ATP synthesis efficiency. Additionally, females may engage compensatory mechanisms to maintain ATP homeostasis without increasing oxidative phosphorylation. Another possibility is that mitochondrial coupling efficiency and proton leak rates differ between sexes, influencing ATP production without affecting overall oxygen consumption. Together, our data support prior research demonstrating that female mitochondria maintain superior bioenergetic efficiency under metabolic stress ^54^. These findings are congruent with the hypothesis of Ganguly, et al. ^58^, that male and female mitochondria adapt/respond to hypoxia (and in the present case, hypothermia) in sex-specific manners, with changes in OXPHOS capacities through different respiratory pathways and utilising different fission/fusion dynamics. Such sex-specific responses may be central to the programmed responses observed in later life.

Given the observed sex differences in growth, cardiovascular morphology and mitochondrial respiration, it is surprising that resting metabolic rate was unaffected by treatment in either of the sexes. Developmental stress is a known risk factor in the development of metabolic syndrome ^62^ and hypoxic incubation has been shown to increase resting 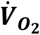 in chicks ^63^. However, in our study, any differences in resting metabolic rates at hatching were resolved by adulthood. Given the differences in heart morphology observed, however, it remains to be seen whether metabolic differences would manifest under stress conditions that place greater demand on the cardiovascular system. For instance, exposure to a treadmill running protocol or acute hypothermia could reveal functional limitations in previously treated males, potentially leading them to reach their 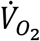 max earlier than controls. Such investigations would be valuable in future studies to assess whether latent cardiovascular deficits become evident under increased physiological strain.

## Conclusions

Together, our data provides evidence that developmental metabolic stress elicits persistent, sex-dependent alterations to cardiovascular function in a vertebrate model in which maternal effects are absent. Our results highlight the necessity of recognizing sex as a crucial determinant in developmental stress and cardiovascular programming. The differential responses observed in males and females underscore the complexity of developmental plasticity, with males exhibiting heightened susceptibility to early-life metabolic stressors in terms of cardiac structural and functional alterations. Despite being evolutionary distant, mammals and birds are both amniotes with convergently evolved endothermy and a four-chambered heart. Indeed, by filling in knowledge gaps regarding developmental programming in birds, we indicate that many cardiovascular responses to developmental stress appear similar between members of these classes regardless of maternal influence. Our findings are therefore broadly relevant not only for birds and reinforce the need for sex-specific approaches in both experimental research and clinical interventions aimed at mitigating the long-term cardiovascular consequences of early-life stress. Advancing our understanding of how developmental stress differentially impacts cardiovascular health across sexes will be pivotal in designing targeted therapeutic strategies that account for these fundamental biological differences and may benefit from the experimental advantages of the avian model.

## Acknowledgements

The authors would like to thank the following funding agencies for their contributions to the research presented: The Swedish Research council (VR) 2019-04053 (CGB, JL). We would also like to thank Monika Hodik and Karin Staxäng for their help in TEM sample preparation and image acquisition. Finally, we would like to thank the staff at SLU Lövsta, in particular Lotta Jönsson and Helena Oscarsson, for their excellent animal care and logistical help.

## Contributions

Experimental design: JL, CGB. Experimental work and sample collection: JL. Conceptualization: JL, CGB. Investigation: JL, SB, BB, CGB, GG, KS. Supervision: CGB. Writing—original draft: JL. Writing—review & editing: JL, CGB, KS, JA, GG.

## Ethical Declarations

Experimental protocols were approved by the Regional Council for Ethical Licensing of Animal Experiments (Uppsala djurförsöksetiska nämnd) and conducted in accordance with all relevant guidelines and regulations (Dnr 5.8.18-18642/2020).

